# Discovery of membrane-permeating cyclic peptides via mRNA display

**DOI:** 10.1101/2020.07.20.212142

**Authors:** John Bowen, Allison E. Schloop, Gregory T. Reeves, Stefano Menegatti, Balaji M. Rao

## Abstract

Small synthetic peptides capable of crossing biological membranes represent valuable tools in cell biology and drug delivery. While several cell-penetrating peptides (CPPs) of natural or synthetic origin have been reported, no peptide is currently known to cross both cytoplasmic and outer embryonic membranes. Here we describe a method to engineer membrane-permeating cyclic peptides (MPPs) with broad permeation activity by screening mRNA display libraries of cyclic peptides against embryos at different developmental stages. The proposed method was demonstrated by identifying peptides capable of permeating *Drosophila melanogaster* (fruit fly) embryos and mammalian cells. The selected peptide cyclo[*Glut*-MRKRHASRRE-*K*\*] showed a strong permeation activity of embryos exposed to minimal permeabilization pretreatment, as well as human embryonic stem cells and a murine fibroblast cell line. Notably, in both embryos and mammalian cells, the cyclic peptide outperformed its linear counterpart and the control MPPs. Confocal microscopy and single cell flow cytometry analysis were utilized to assess the degree of permeation both qualitatively and quantitatively. These MPPs have potential application in studying and non-disruptively controlling intracellular or intraembryonic processes.

## Introduction

Cell- and embryo-permeating technologies are valuable tools in the study of intracellular processes in biology.^1,2^ A number of chemical and physical methods are available for introducing molecular probes into single cells and embryos to investigate intracellular metabolic and signaling pathways.^2–8^ Among physical tools, microinjection is widely employed to access the complex architecture of embryos.^1,2^ Among chemical tools, peptides have received considerable attention. A large number of cell-permeating peptides (CPPs) with either broad or cell-specific permeating activity have been previously described.^4,9–12^ Peptides have several characteristics that make them preferred delivery vectors compared to physical tools. In particular, they provide excellent delivery rates while causing minimal or no disruption of the cell membrane and intracellular structures.^13^ While largely available for intracellular delivery, however, peptide vectors capable of accessing embryos are not known. To overcome this limitation, we sought to develop membrane-permeating peptides (MPPs) that enable access to both embryos and cells, using a combinatorial screening approach.

The development of CPPs follows two routes, namely rational design or combinatorial screening. In the context of embryo permeation, rational design is not a viable route, given the lack of detailed information on the biomolecular requirements (*e*.*g*., molecular weight cut-off, charge and amphiphilicity) needed to overcome the complex, multilayer barrier of embryos. Therefore, we undertook the route of screening combinatorial libraries of cyclic peptides, constructed via mRNA display technology. In mRNA display, a peptide is covalently linked to its corresponding mRNA transcript via a puromycin translation inhibitor and psoralen DNA intercalating agent.^14^ The mRNA is stabilized by its complementary DNA and purified via orthogonal purification techniques that leverage the poly adenylated tail and the oligo-histidine tag integrated in the mRNA-peptide hybrid. Previous studies have reported the use of mRNA display to identify linear cell-permeating peptides.^10^ Cyclic peptides, however, are particularly attractive as cell permeating peptides (CPPs) due to their conformational-induced resistance to proteolytic enzymes, which prolongs their activity in the peri-/intra-cellular milieu. Recent efforts on the discovery of novel permeating peptides include the use of a phylomer peptide library;^15^ others have identified short peptide motif sequences as a general vehicle of intracellular delivery of cyclic peptides,^16^ and subsequently modified the sequence and/or stereochemistry to enhance its activity.^17^ Pei and other researchers have discovered that cyclization of arginine rich sequences increases cellular uptake and efficiency.^16–21^ Nevertheless, collectively the discovery and characterization of cyclic CPPs represents only 5% of all the experimentally derived sequences published on the CPP site 2.0 database.^11^

Our group has developed a method to conduct peptide cyclization upon adsorbing the hybrids onto an oligo dT solid phase, which employs amine-reactive crosslinkers to tether the peptide N-terminus with the C-terminal side chain ε-amino group of lysine.^22^ To identify cyclic MPPs, we screened mRNA display libraries of cyclic peptides against *D. melanogaster* embryos and identified candidate embryo-permeating sequences. We then evaluated a selected candidate sequence in both its linear and cyclic forms, to penetrate *D. melanogaster* embryos as well as NIH 3T3 cells and H9 human embryonic stem cells (hESCs). Cationic (*i*.*e*., the known cell-permeating peptide poly-arginine) and anionic peptides were employed as positive and negative controls, respectively. The broad membrane-permeating activity of the selected peptide was confirmed by imaging the embryos and cells via laser-scanning confocal fluorescence microscopy, and by analyzing the cells via flow cytometry. Notably, the selected cyclic peptide cyclo[*Glut*-MRKRHASRRE-*K*\*] was found to permeate both embryos and cells more efficiently compared to both its linear counterpart and the control sequences, indicating that both amino acid sequence and cyclization are key to endow membrane-permeating activity.

## Materials and Methods

### Production of mRNA display libraries of cyclic 10-mer peptides

An mRNA-display library of cyclic 10-mer peptides was initially produced following the method described by Menegatti *et al*.^22^ All oligonucleotides were obtained from Integrated DNA Technologies (IDT). First, the oligonucleotide sequence 5’-GGA CAA TTA CTA TTT ACA ATT ACA ATG *<u>NNN NNN NNN NNN NNN NNN NNN NNN NNN NNN</u>* AAA GGC GGC AGC GGC GGC AGC CAT CAC CAC CAT CAC CAT ATG GGA ATG-3’ encoding the 10-mer (MX_1_X_2_X_3_X_4_X_5_X_6_X_7_X_8_X_9_X_10_K) library (where X is one of the 20 natural amino acids) was PCR-amplified with Phusion DNA polymerase (Fisher Scientific), using the forward primer 5’-GCA AAT TTC TAA TAC GAC TCA CTA TAG GGA CAA TTA CTA TTT ACA ATT AC-3’ and the reverse primer 5’-ATA GCC GGT GCC AGA TCC AGA CAT TCC CAT ATG GTG ATG GT-3’. The amplified DNA was used to generate mRNA-peptide fusions using *in vitro* transcription using retic lysate IVT Kit (Fisher Scientific), ligation of a puromycin hybridization and intercalation linker (Keck Oligo Synthesis Lab, Yale University), and *in vitro* translation. The resulting mRNA-peptide libraries (∼10^13^ hybrids) were adsorbed onto 200 µL of Oligo-dT cellulose (Fisher Scientific), washed with dT washing buffer (0.2 M NaCl, 10 mM Tris pH 7.5, 1 mM EDTA, 0.05% SDS, 1 mM DTT (Fisher Scientific) 3 × 10 min) and crosslinking buffer (0.2 M NaCl, 1 mM EDTA, 0.05% SDS, 1 mM DTT; 3 × 10 min), and finally cyclized by incubation with 50 µL of 2.3 mg/mL solution of DSG (Fisher Scientific) in DMF (Fisher Scientific) for 2 hrs at 4°C. The mRNA-cyclic peptide libraries were eluted with 600 µL of 0.1% DEPC (Millipore Sigma) water containing 1 mM DTT. Next, the libraries were incubated with 25 µL of 100 µM reverse transcription primer (5’-TTT TTT TTT TNN CCA GAT CCA GAC ATT CCC AT-3’) for 15 min at room temperature, 200 µL of 5x first strand buffer (250 mM Tris–HCl pH 8.3, 375 mM KCl, 15 mM MgCl_2_), and 50 µL of 10 mM dNTPs (New England BioLabs), 100 µL 0.1 M DTT, and 0.1% DEPC water to adjust the solution to a total volume to 995 µL. Reverse transcription was initiated by incubating with 5 µL Superscript™ reverse transcriptase (200 U/µL) (Fisher Scientific) for 50 min at 42°C; EDTA was added to the reaction mixture to reach a final concentration of 6mM and incubated for 5 min at room temperature. The resulting libraries of cDNA-mRNA-cyclic peptides were desalted using a Nap-10 desalting column and purified using Ni-NTA agarose beads. The purified libraries were finally desalted using a Nap-5 column. The estimated number of mRNA-cDNA-cyclic peptide fusions, determined by *A*_260_ measurement, was ∼10^13^ variants.

### Screening of mRNA cyclic peptide libraries against D. melanogaster embryos

Adult *Drosophila melanogaster* flies (∼100 units) were incubated for 2-4 hrs in a cage with an agar plate containing grape juice and yeast extract (3% w/v agar, 25% grape juice concentrate, 0.03% methyl paraben (p-hydroxymethylbenzoate), and 0.3% w/v sucrose) (Genesee Scientific). The flies were anesthetized with CO_2_ for 3 – 5 sec, after which the existing agar plate was replaced with a fresh plate, and flies were allowed to deposit embryos. The plate was removed from the cage after 2 hrs and 50-100 embryos were extracted, placed onto a mesh screen, washed with embryo wash buffer (3.5 M NaCl, 1.5% v/v Triton X-100 in water), dechorionated with 8.25% sodium hypochlorite in water for 3 min and finally washed with PBS (Millipore Sigma). The embryos were suspended and washed in a microcentrifuge tube containing 1 mL of PBS, pH 7.4, and finally settled by gravity. After removing the PBS supernatant, the entirety (∼1 mL) of mRNA-cDNA-cyclic 10-mer peptides was added to the embryos and incubated at room temperature for 3 hrs under mild agitation. The embryos were washed with PBS buffer and lysed using a shortened version of TRIzol (Fisher Scientific) lysis method; briefly, the method provided by the manufacturer was followed up to the partitioning step. A volume of 1 mL TRIzol lysis buffer was incubated with the embryos, and a 21-gauge needle was used to lyse the embryos. The lysed embryos were centrifuged at 12,000x g for 20 min to pellet any membrane components or un-lysed embryos. The mRNA-cDNA-cyclic peptides were precipitated using 2 mL of 100% v/v ethanol and 100 µL of 3 M Potassium Acetate pH 5.5 overnight at −20°C. The pellet was centrifuged at 12,000x g for 20 minutes and washed with 70%v/v and 100%v/v ethanol in series, and finally resuspended in 30 µL of autoclaved 0.1% DEPC water. The cDNA was amplified by PCR as described above and utilized as a template to construct a subsequent iteration of the mRNA-display library. Four subsequent cycles of library screening were repeated, and the final pool of cDNA sequences were amplified by PCR, and sequenced by Sanger sequencing.

### Culturing of H9 human embryonic stem cells and NIH 3T3 cells

H9 hESCs were initially seeded as colonies on 3.5 cm^2^ vitronectin-coated polystyrene tissue culture plates (Sarstedt) and expanded in E8^®^ media (Stemcell Technologies). Cells were cultured and passaged upon reaching approximately 70% confluency. Cells were washed with PBS, released from the plates using 1 mL of ReLeSR^®^ (Stemcell Technologies), and resuspended by mechanical agitation. Finally, 50 µL of cell suspension was added to a fresh vitronectin-coated plate, and allowed to grow for 5 - 6 days until reaching approximately 70% confluency, while replenishing with fresh E8^®^ media every 1-2 days. NIH 3T3 cells were initially seeded on 3.5 cm^2^ polystyrene tissue culture plates and expanded in DMEM^®^ + GlutaMAX™ (Gibco), supplemented with 10%v/v One Shot™ Fetal Bovine Serum (Gibco) and 100 U/mL penicillin-streptomycin (Gibco). Cells were passaged upon reaching approximately 70% confluency, the cells were washed with PBS and released from the plates using 500 µL of TrypLE™ (Gibco), incubated at 37°C and centrifuged at 300x g for 5 min. Subsequently, the supernatant was removed and the cell pellet was resuspended in 1 mL of DMEM^®^ + GlutaMAX™. Finally, the cells were passed at a 1:10 split ratio and allowed to grow for 3 - 4 days until reaching approximately 70% confluency, while changing the media every 2 days.

### Evaluation of peptide permeation on D. melanogaster embryos and larvae

*D. melanogaster* embryos at different developmental stages and larvae were allowed to propagate using grape juice agar plates, yeast extract, and mesh cages as described in previous sections. A cage of fruit flies were allowed to propagate and lay embryos for 48 hours onto the grape juice agar plate. The plate was subsequently collected, the embryos/larvae were removed from the plate by washing with 3 × 30 mL of embryo wash buffer (3.5 M NaCl, 1.5% v/v Triton X-100 in water), and placed on a mesh screen. The embryos/larvae were subsequently treated with 10 mL of 8.25% sodium hypochlorite in water for 3 min to remove the embryonic chorion, and washed with PBS prior to peptide incubation. After dechorionation, a fraction of the embryos and larvae were also treated with 10%v/v of detergent Valencia Orange Concentrated Degreaser (Citrasolv) in water for 3 minutes, which dewaxes the embryos, as described by Rand *et al*.^23^ Dechorionated and dechorionated/dewaxed embryos/larvae were washed and resuspended in 500 µL of MBIM media (MgCl_2_·6H_2_0 (2.2 g/L), MgSO_4_·7H_2_0 (2.97 g/L), NaH_2_PO_4_ (0.42 g/L), Glutamic Acid (12.1 g/L), Glycine (6.05 g/L), Malic Acid (0.66 g/L), Sodium Acetate (0.027 g/L), Glucose (2.2 g/L), CaCl_2_·2H_2_0 (0.99 g/L)) supplemented with 0.05% Tween 20 (Promega), 2.5 mM DTT, at pH 7.4.

Aqueous stock solution of peptides cyclo[*Glut*-MRKRHASRRE-*K*\*]-K(5-TAMRA) (UNC Peptide Core Facility), TAMRA-MRKRHASRREK, TAMRA-RRRRRRRRR, and TAMRA-GEGEGEG (Genscript) at 0.1 mM concentration were added to the suspension of embryos/larvae to reach a final concentration of 20 µM and incubated overnight at room temperature under gentle agitation; a control with no peptide (PBS, pH 7.4) was also performed. These trials resulted in a pool of embryos and larvae featuring development ages ranging between 12 – 60 hours, which includes the 48 hour fruit fly propagation period and the ∼12 hour overnight incubation period. The embryos/larvae were washed 3x with 1 mL of MBIM, 0.05% Tween 20, 2.5 mM DTT, and analyzed by laser scanning confocal microscopy in the mid-sagittal section using a Zeiss LSM 880 with Airyscan instrument (White Plains, New York) operating at varying excitation wavelengths. An excitation wavelength of 561 nm with an emission filter range of 570 – 650 nm was used for TAMRA detection in the red channel. An excitation wavelength of 488 nm and emission filter range of 490 – 570 nm was used to detect GFP expression within the embryo nuclei in the green channel. Image analysis of nuclei localization and peptide distribution was performed via Zeiss image analysis software.

### Evaluation of peptide permeation on H9 hESCs and NIH 3T3 cells

H9 hESCs were initially seeded on a 3.5 cm^2^ vitronectin-coated plate and allowed to propagate as colonies as described in previous sections. Cells were allowed to reach ∼70% confluency, released from the plate using TrypLE^®^, resuspended in 1 mL of E8^®^ media, centrifuged at 300x g for 5 min, and plated at 50,000 cells per plate in 2 mL of E8^®^ media supplemented with 2 µL of 1000x Rho/Rock inhibitor Y27632 (Tocris). Prior to incubation with the appropriate peptide, plated cells were allowed to grow for 48 hrs and washed with PBS, after which 600 µL of fresh E8^®^ media either pre-warmed to 37°C or pre-chilled to 4°C was added to the tissue culture dish. Aqueous solutions of peptides cyclo[*Glut*-MRKRHASRRE-*K*\*]-K(5-TAMRA), TAMRA-MRKRHASRREK, TAMRA-RRRRRRRRR, and TAMRA-GEGEGEG at 0.1 mM concentration were added to reach a final concentration ranging from 1 – 10 µM in E88^®^ media. Cells were incubated with the peptides for 3 hrs at either 37°C in 5% CO_2_ or at 4°C, and then washed 2x with 1 mL PBS supplemented with MgCl_2_ and CaCl_2_ (Millipore Sigma), 1x with 1 mL of 20 mM glycine, 3 M urea, at pH 3.0 for 5 minutes at 4°C, and 2x with 1 mL PBS added with supplemented with MgCl_2_ and CaCl_2_.

NIH 3T3 cells were cultured following the procedure outlined above. Cells were allowed to reach ∼70% confluency, released from the plate using TrypLE^®^, resuspended in 1 mL of DMEM^®^ + GlutaMAX™, supplemented with 10%v/v One Shot™ Fetal Bovine Serum and 100 U/mL penicillin-streptomycin, centrifuged at 300x g for 5 min, and plated at 50,000 cells per plate in 2 mL of DMEM^®^ + GlutaMAX™, supplemented with 10%v/v One Shot™ Fetal Bovine Serum and 100 U/mL penicillin-streptomycin. Prior to incubation with peptides, plated cells were allowed to grow for 48 hrs and washed with PBS, after which 600 µL of fresh in Opti-Mem® media either pre-warmed to 37°C or pre-chilled to 4°C was added to the tissue culture dish. Aqueous solutions of peptides cyclo[*Glut*-MRKRHASRRE-*K*\*]-K(5-TAMRA), TAMRA-MRKRHASRREK, TAMRA-RRRRRRRRR, and TAMRA-GEGEGEG at 0.1 mM concentration were added to reach a final concentration of 1 – 10 µM in Opti-Mem^®^ media. Cells were incubated with the peptides for 3 hrs at either 37°C in 5% CO_2_ or at 4°C, and then washed 2x with 1 mL PBS supplemented with MgCl_2_ and CaCl_2_, 1x with 1 mL of 20 mM glycine, 3 M urea, at pH 3.0 for 5 minutes at 4°C, and 2x with 1 mL PBS supplemented with MgCl_2_ and CaCl_2_.

For single cell flow cytometry analysis, following the washing steps, both H9 hESCs and NIH 3T3 cells were removed from the plate using 500 µL of TrypLE^®^ and centrifuged at 300x g for 5 min, after which the supernatant was removed and the cell pellet was resuspended in 200 µL of PBS. Suspended cells were then analyzed using a MACS Quant Flow Cytometer (Miltenyi Biotec) operated at the excitation wavelength of 561 nm and with an emission filter of 661/20 nm. A total of 10,000 events were recorded.

For analysis using laser scanning confocal microscopy, both H9 hESCs and NIH 3T3 cells were grown on 3.5 cm^2^ glass-bottom tissue culture plates (Grenier-Bio-One), stained with either Hoechst nuclear dye alone (1 µg/mL in PBS, pH 7.4) followed by Hoechst nuclear dye, and incubated on ice for 15 minutes. Following a wash with PBS supplemented with MgCl_2_ and CaCl_2_, cells were imaged by laser scanning confocal microscopy using a Zeiss LSM 880 with Airyscan instrument (White Plains, New York) operated at the excitation wavelength of 561 nm with an emission filter of 570 - 610 nm to detect TAMRA expression in the red channel. An excitation wavelength of 405 nm and emission filter of 410 – 490 nm was used to detect Hoechst dye indicating nuclear localization. Image analysis of peptide distribution was performed via Zeiss image analysis software.

### Evaluation of cytotoxicity of cyclo[Glut-MRKRHASRRE-K*]-K(5-TAMRA) on H9 hESC and NIH 3T3 cells

H9 hESCs and NIH 3T3 cells were cultured as described above. Aqueous stock solutions of peptides cyclo[*Glut*-MRKRHASRRE-*K*\*]-K(5-TAMRA), TAMRA-RRRRRRRRR, and TAMRA-MRKRHASRREK at 0.1 mM concentration were added to the cell culture solutions to reach a final concentration of 10 µM in either E8^®^ media or DMEM^®^ + GlutaMAX™, supplemented with 10%v/v One Shot™ Fetal Bovine Serum and 100 U/mL penicillin-streptomycin for H9 hESCs or NIH 3T3 cells, respectively. Cells were incubated with the peptides for 24 or 48 hrs at 37°C, 5% CO_2_, subsequently washed twice with PBS, released from the tissue culture plate using TrypLE^®^, centrifuged at 300x g for 5 min and resuspended in 1 mL PBS. Cell viability was determined using a Live/Dead™ Fixable Dead Cell Stain Kit. Briefly, the cells were incubated with Live/Dead™ Fixable Green Dead Cell Stain dye (Fisher Scientific) according to the manufacture’s protocol, and washed twice with PBS, 1% BSA prior to flow cytometry analysis. Suspended cells were then analyzed using a MACS Quant Flow Cytometer (Miltenyi Biotec) operated at the excitation wavelength of 561 nm and emission filter of 661/20 nm for detection of the TAMRA fluorophore, and operated at an excitation wavelength of 488 nm and emission of 525/50 nm for detection of the FITC dye. A total of 10,000 events were recorded.

## Results and Discussion

### Identification of candidate D. melanogaster embryo-permeating peptides via screening of mRNA display cyclic peptide libraries

The complex barrier that envelops developing fruit fly embryos comprises an exochorion (300-350 nm), an endochorion (500-700 nm), an innermost chorion (40-50 nm), a thin waxy layer (∼5 nm)^24^, and a vitelline membrane (300 nm).^25^ The delivery of bioactive compounds such as DNA-nanoparticle complexes^1^, Cas9-RNA complexes^26^, or double stranded RNA^27^ into developing fruit fly embryos is commonly performed using microinjection. While effective, this technology is limited by its inherently low throughput, which poses the need for liquid-phase delivery vectors that can reliably deliver a payload into a large number of embryos. Organic solvents such as heptane, octane, toluene, ethanol, or monoterpenes such as D-limonene have been shown to permeabilize developing fruit fly embryo^23,24,28^ and developing wheat germ embryos.^29^ These solvents, however, are toxic and can cause irreparable damage to the embryo membrane. VisuFect, a proprietary organic compound, has been recently developed to permeate into the embryos of multiple organisms and deliver cargo;^6,30^ currently, however, no data on the composition of VisuFect, its mechanism of permeation, and potential adverse effect on embryos are available.

Advanced molecular delivery vectors^31^ (*e*.*g*., small organic molecules, peptides, and proteins) are not available for embryos, despite their success with single cells.^5,8,32–38^ Peptides are particularly attractive for this application: they can be produced at large volumes affordably, are safety and biocompatibility, and can act as both penetrating agents and probes for manipulating PPIs and NPIs.^39^ The screening of peptide libraries has been shown to be a powerful tool for the identification of cell- and tissue-permeating peptides.^40^ The underlying premise of combinatorial screening is that a peptide sequence possessing any desired bioactivity can be identified, provided that the library has high diversity, and the screening process is appropriately designed and conducted through a sufficient number of iterations.

Accordingly, we constructed an mRNA display library of cyclic 10-mer peptides and developed a method of screening against dechorionated *D. melanogaster* embryos to identify embryo-permeating peptides. The embryo manipulation and screening procedure is shown in **Figure S1**: *(i)* following an initial growth on yeast extract-supplemented agar plates, the embryos are collected, washed, and dechorionated/dewaxed; *(ii)* the prepared embryos are incubated with the library; *(iii)* following incubation, the embryos are lysed, and the library-associated cDNA is amplified via PCR and utilized to construct subsequent libraries; starting at the second round of screening, a membrane/organelle centrifugation step is added after the lysis of the embryos to avoid the selection of peptide sequences that are either adsorbed to the vitelline membrane or absorbed within the embryo vitelline membrane but do not possess a true permeation activity (*i*.*e*., false positives). Four rounds of screening were performed in this study to ensure the selection of peptides with high permeation activity, resulting in the sequences listed in **Table 1**. Of the 17 identified sequences, one 9-mer (Sequence 11) and six 10-mer (Sequences 12-17) peptides were considered as leads owing to their high chemical diversity. The presence of contaminant pentameric and heptameric sequences resulting from mRNA splicing and degradation, and truncated sequences resulting from sequencing errors (nonamers) may be due to contamination in the oligonucleotide pool used to generate the initial library.

**Table 1.** Sequences identified by screening mRNA display libraries of cyclic peptides against dechorionated *D. melanogaster* embryos.

|  | M | X<sub>1</sub> | X<sub>2</sub> | X<sub>3</sub> | X<sub>4</sub> | X<sub>5</sub> | X<sub>6</sub> | X<sub>7</sub> | X<sub>8</sub> | X<sub>9</sub> | X<sub>10</sub> | K |
| --- | --- | --- | --- | --- | --- | --- | --- | --- | --- | --- | --- | --- |
| **EPP.1** |  | C | T | C | F | T |  |  |  |  |  |  |
| **EPP.2** |  | Y | K | C | N | L |  |  |  |  |  |  |
| **EPP.3** |  | A | A | N | C | S |  |  |  |  |  |  |
| **EPP.4** |  | G | L | R | T | P |  |  |  |  |  |  |
| **EPP.5** |  | S | K | E | R | S | F | V |  |  |  |  |
| **EPP.6** |  | S | K | E | T | S | F | V |  |  |  |  |
| **EPP.7** |  | L | R | N | L | Q | S | K |  |  |  |  |
| **EPP.8** |  | L | V | W | Q | G | F | R |  |  |  |  |
| **EPP.9** |  | S | K | E | T | S | F | V |  |  |  |  |
| **EPP.10** |  | S | K | E | T | S | F | V |  |  |  |  |
| **EPP.11** |  | R | K | R | H | A | S | R | R | E |  |  |
| **EPP.12** |  | S | V | N | L | L | R | C | E | A | R |  |
| **EPP.13** |  | C | I | Y | L | K | I | M | S | L | I |  |
| **EPP.14** |  | W | N | V | G | L | D | H | V | R | G |  |
| **EPP.15** |  | G | E | V | M | G | M | S | L | L | I |  |
| **EPP.16** |  | W | S | R | S | V | R | E | A | V | S |  |
| **EPP.17** |  | D | T | P | E | R | R | S | A | R | G |  |

Sequence analysis shows a clear enrichment in positively charged amino acids, especially arginine (R), against a rather marginal enrichment of negatively charged amino acids. A mild enrichment in hydrophobic, non-aromatic amino acids (leucine, isoleucine, valine, alanine, and methionine), and the accumulation of aromatic amino acids (tryptophan and tyrosine) on the sequence N-terminus is also observed. The enrichment is cationic, specifically arginine residues, and aliphatic/aromatic amino acids is known to endow peptides with the ability to cross biological barriers.^17,21,41^ At the same time, the presence of hydrogen-binding amino acids (serine), together with the cationic residues, endows the selected peptides with a multipolar character and good water solubility.

Several pentamer (sequences 1-3) and decamer sequences (sequences 12 and 13) feature the presence of cysteine residues, which can form disulfide bonds with free cysteines present on the surface of proteins. Relevantly, the vitelline membrane in *D. melanogaster* embryos is rich in cysteine and features a dense disulfide-crosslinked network, particularly in the later stages of development.^42^ We therefore speculate that the selection of cysteine-containing sequences, especially among pentamers, was driven by the formation of disulfide bonds between the library peptides and the vitelline membrane of the embryos during screening. On the other hand, the presence of sulfur-containing amino acids in bioactive peptides is not desired, due to their tendency to form undesired disulfide bonds (cysteine) or be oxidized to their sulfone derivatives (methionine), which alters irreversibly their biochemical activity. Accordingly, these sequences were not considered for experimental evaluation. Further, decamer sequence 14 was predicted to have poor water solubility and was also discarded. Of particular interest is the nonamer sequence 11, which features a calculated isoelectric point of 12.6 and a net charge of +5.1 at pH 7, high water solubility, and multipolar character. Notably, peptide RRRRRRRRR (poly-arginine), a known cell- and tissue-permeating peptide, is also a nonamer. Based on these considerations, sequence 11 was selected for characterization via embryo- and cell-permeating tests.

### Qualitative assessment and ranking of Drosophila melanogaster permeability, viability, and developmental stage

The permeation of the selected cyclic peptide cyclo[*Glut*-MRKRHASRRE-*K*\*] and its linear counterpart MRKRHASRREK in *D. melanogaster* was evaluated via fluorescence confocal microscopy; since peptides known to permeate *D. melanogaster* embryos are not available, poly-Arg, a known cationic cell-permeating peptide, was adopted as positive control, while anionic peptide GEGEGEG was designed as a negative control. All peptides were labeled with a rhoda-mine-derived fluorophore TAMRA (the cyclo[*Glut*-MRKRHASRRE-*K*\*]-K(5-TAMRA) is reported in **Figure S2**).

Previous studies have demonstrated that successful embryo permeation requires the removal of the chorionic barrier.^23^ The contiguous vitelline membrane and wax layer, underlying the chorionic barrier, also pose a challenging barrier due to their connected, dense, and irregular structures.^24,25^ To parse the resistance to peptide permeation opposed by the chorionic barrier and the vitelline/wax barrier, we resolved to test both dechorionated-only and dechorionated-and-dewaxed embryos. The pretreated embryos were incubated with 20 µM of either the test or control TAMRA-labeled peptides overnight at room temperature, and subsequently imaged by confocal fluorescence microscopy (*note:* Tween-20, a mild surfactant, and DTT were added to the peptide solutions to avoid non-specific hydrophobic adsorption and entrapment of the peptides on the highly disulfide-crosslinked vitelline membrane).^42^ To account for the differential permeability of the various layers, peptide adsorption onto the membrane and permeation in the interior of the *D. melanogaster* were evaluated separately, the former as a thin, outer layer of red fluorescence, and the latter as diffuse/punctate red fluorescence.

Furthermore, we evaluated both the viability and developmental stage (*i*.*e*., embryo *vs*. larva) of the embryos used in this study, together with the peptide-induced fluorescence on their surface and interior. Specifically, we adopted two criteria, namely the auto-fluorescence of the developing yolk in the blue channel, which has been associated with the health of the embryos,^23^ and the expression of green fluorescent protein (GFP), which reveals whether the host is at the embryonal or larval state based on the location of the nuclei and the structure of the membrane. While collected as embryos, in fact, the hosts were at different stages of early development when incubated with the peptides. Due to the long incubation time needed for the peptides to permeate the hosts effectively (12 hrs), several embryos transitioned to larvae or showed a loss in viability. To account for such variability and ensure statistical significance of our observations we performed a total of 96 peptide incubations across dechorionated-only and dechorionated-and-dewaxed hosts. The results of the qualitative assessment of the collected images are summarized in **Figure 1** for the dechorionated-only hosts and in **Figure 2** for the dechorionated-and-dewaxed hosts.

**Figure 1.**
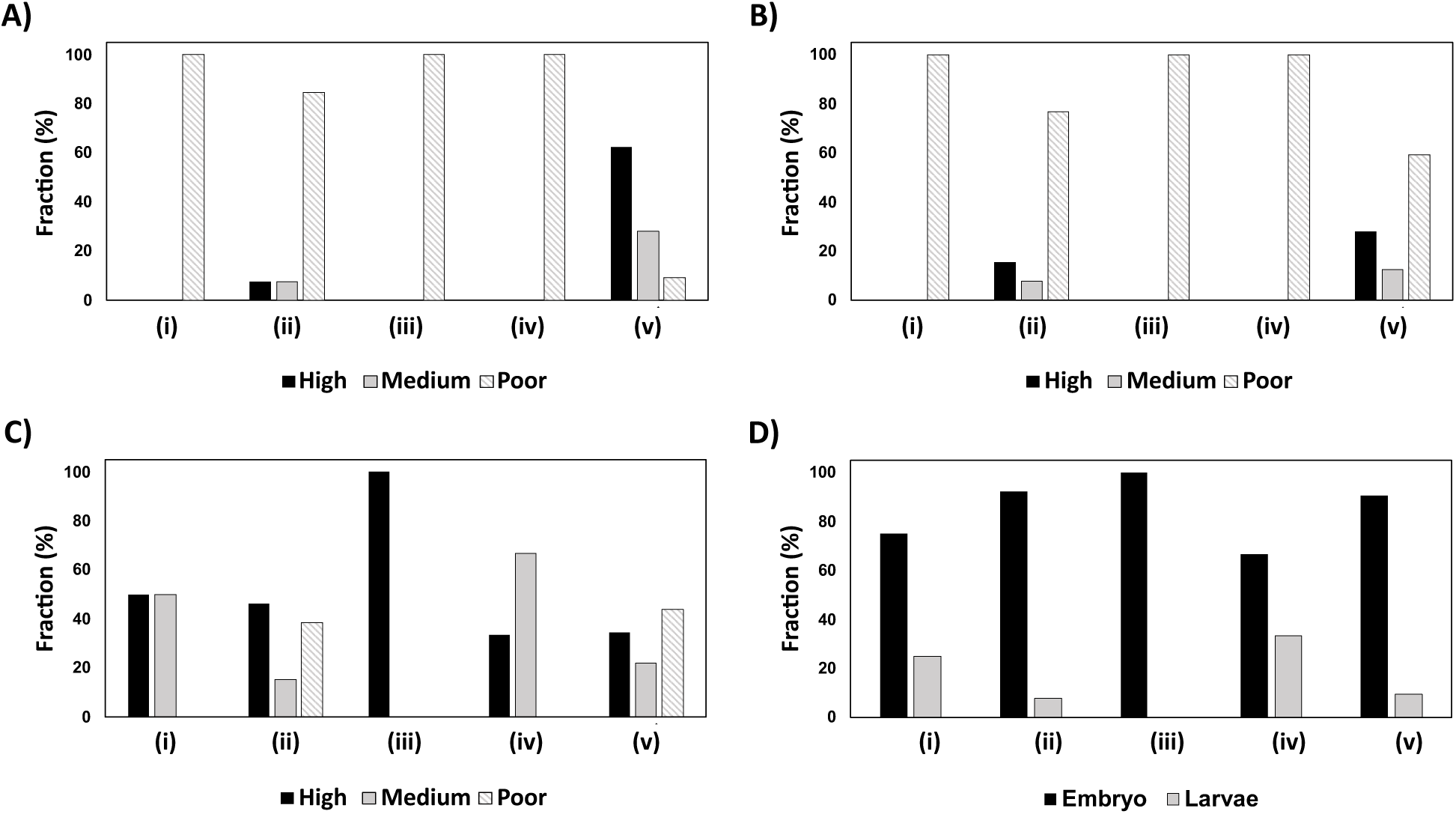
Qualitative assessment of peptide **(A)** adsorption on the outer membrane and **(B)** permeation in the interior, together with **(C)** viability and **(D)** developmental stage of dechorionated *Drosophila melanogaster* hosts incubated with either *(i)* no peptide (PBS, pH 7.4), *(ii)* GEGEGEG, *(iii)* RRRRRRRRR, *(iv)* MRKRHASRREK, or *(v)* cyclo[*Glut*-MRKRHASRRE-*K*\*] peptides. Peptide adsorption onto the membrane was assessed as a thin layer of red fluorescence, while peptide permeation in the interior was assessed as diffuse or punctate red fluorescence within the developing embryo or larva. The level of superficial and inner fluorescence (**A** and **B**), as well as host viability **(C)** were qualitatively categorized as either “high”, “medium”, or “poor”. Accordingly, the axis “fraction (%)” indicates the fraction of the hosts tested under conditions *(i) – (v)* that belonged to either the “high”, “medium”, or “poor” categories.

**Figure 2.**
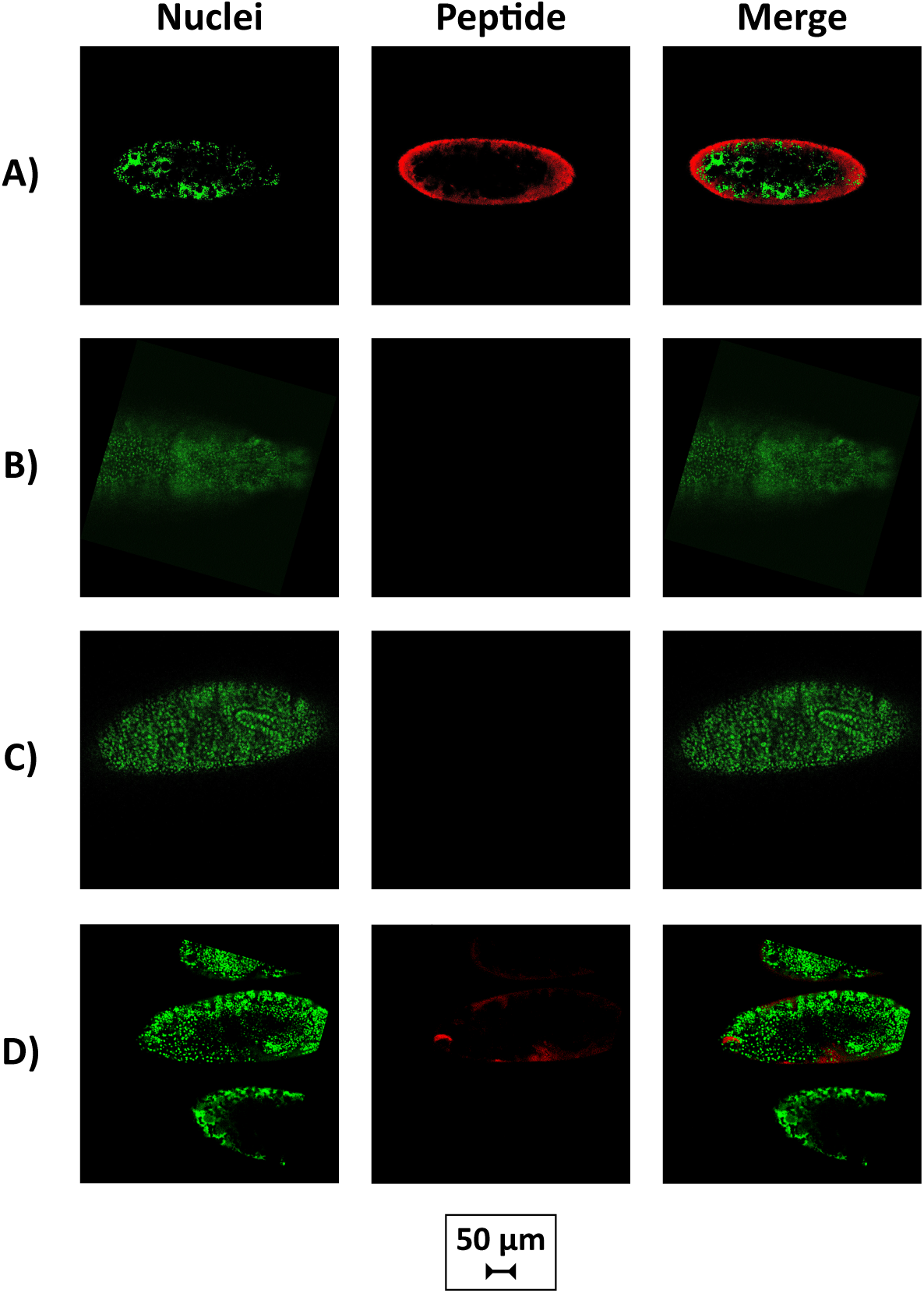
Confocal fluorescence microscopy images of dechorionated *D. melanogaster* embryos or larvae incubated with TAMRA-labeled peptides **(A)** cyclo[*Glut*-MRKRHASRRE-*K*\*], **(B)** MRKRHASRREK, **(C)** RRRRRRRRR, and **(D)** GEGEGEG at 20 μM, overnight at room temperature (*note:* the scalebar is reported separately from the microscopy images to improve its legibility). Nuclear GFP expression is shown in the green channel, and distribution of permeated peptides in the hosts is shown in the red channel.

Peptide cyclo[*Glut*-MRKRHASRRE-*K*\*]-K(5-TAMRA) was found to adsorb onto and permeates into dechorionated-only hosts (**Figure 1A** and **1B**) to a greater extent than all the other peptides. As anticipated, negative control GEGEGEG and the linear cognate MRKRHASRREK featured poor or no permeation on all tested embryos/larvae. Somewhat unexpected was the behavior of RRRRRRRRR, a prominent cell-permeating peptide, which was found not to adsorb or permeate the hosts. These results suggest that the physicochemical properties that make a peptide capable of permeating cells differ from those required to endow a peptide with embryo-permeating activity, and highlight the effectiveness of cyclo[*Glut*-MRKRHASRRE-*K*\*] as an embryo-permeating vector.

A partial loss of viability and a transition to larval state were observed across all the tested embryos, with the sole exception of those treated with RRRRRRRRR. On the other hand, loss of viability (**Figure 1C**) and conversion to larvae (**Figure 1C**) occurred respectively in 50% and 25% of the embryos treated with no peptide (PBS, pH 7.4); similarly, a loss of viability of 60% and 30% conversion to larvae was observed with the linear cognate MRKRHASRREK, which showed completely null adsorption onto and permeation into the hosts. We therefore assume this to be an inherent behavior of the hosts and not a result of the contact with the peptides. At the same time, neither the viability level nor the developmental status seemed to impact peptide permeation, since comparable levels of superficial and inner fluorescence were observed in embryos and larvae alike.

Confocal microscopy images exemplifying these results are reported in **Figure 2**: specifically, **Figure 2A** shows the permeation of the lead cyclic peptide cyclo[*Glut*-MRKRHASRRE-*K*\*]-K(5-TAMRA) in a dechorionated *D. melanogaster* embryo; together with permeation, a considerable localization of the peptide can be observed on the barrier formed by the vitelline membrane coated by the wax layer. The latter is a likely substrate for hydrophobically driven adsorption of small organic molecules like peptides. On the other hand, the linear precursor TAMRA-MRKRHASRREK (**Figure 2B**) did not show any accumulation at this layer, and neither did the control peptides TAMRA-RRRRRRRRR (**Figure 2C**) and TAMRA-GEGEGEG (**Figure 2D**). The lack of interaction between MRKRHASRREK and the vitelline/wax barrier suggests that cyclo[*Glut*-MRKRHASRRE-*K*\*] does not adsorb irreversibly on the wax layer, but it rather permeates through it to reach the underlying vitelline membrane, through which it permeates, ultimately accessing the embryo’s interior. At the same time, the poor permeability of MRKRHASRREK speaks to the importance of peptide cyclization towards permeation activity; as indicated in **Figure 1D**, upon incubation with MRKRHASRREK, a substantial fraction of the embryos (30%) transitioned to larvae, of which one specimen is shown in **Figure 2B**. Finally, a representative three dimensional z-stack analysis of a cyclo[*Glut*-MRKRHASRRE-*K*\*]-K(5-TAMRA) incubation, shown in **Figure S3**, demonstrates the permeation of the cyclic peptide through the interior of the embryo. Most importantly, the diffuse fluorescence around the perimeter of the embryo diffusing toward the interior confirms that the cyclic peptides are not blocked by adsorption on the peripheral vitelline membrane and permeate through the entirety of the embryo.

We then sought to investigate peptide permeation in dechorionated-and-dewaxed *D. melanogaster* hosts (**Figure 3**). As the screening was conducted on embryos carrying an intact wax layer, we hypothesized that its removal would enhance the peptide permeation flux. The wax layer consists of unsaturated fatty acids and paraffins, and acts as a “water-proofing” layer preventing inner embryo from desiccation.^23^ The removal of the wax layer using organic solvents (“dewaxing”) has been shown to promote permeation of small organic compounds such as cycloheximide, methylmercury chloride, cytochalasin D, cyclophosphamide, and Nocodazole.^23,28^ The combination of dechorionation and dewaxing maintains the vitelline membrane, a largely proteinaceous membrane mostly comprising serine, proline, glycine, and alanine residues, as the only layer enveloping the embryo and yolk.^43^ Following dechorionation, the embryos were dewaxed using 10%v/v Citrasolv in MBIM buffer, washed, and incubated with the fluorescently labeled peptides (*note:* another dewaxing solvent has also been described for permeabilizing *D. melanogaster* embryos to small molecules and fluorescent dyes^28^, which comprises a combination of limonene, heptane, monoterpene oil, and surfactants).

**Figure 3.**
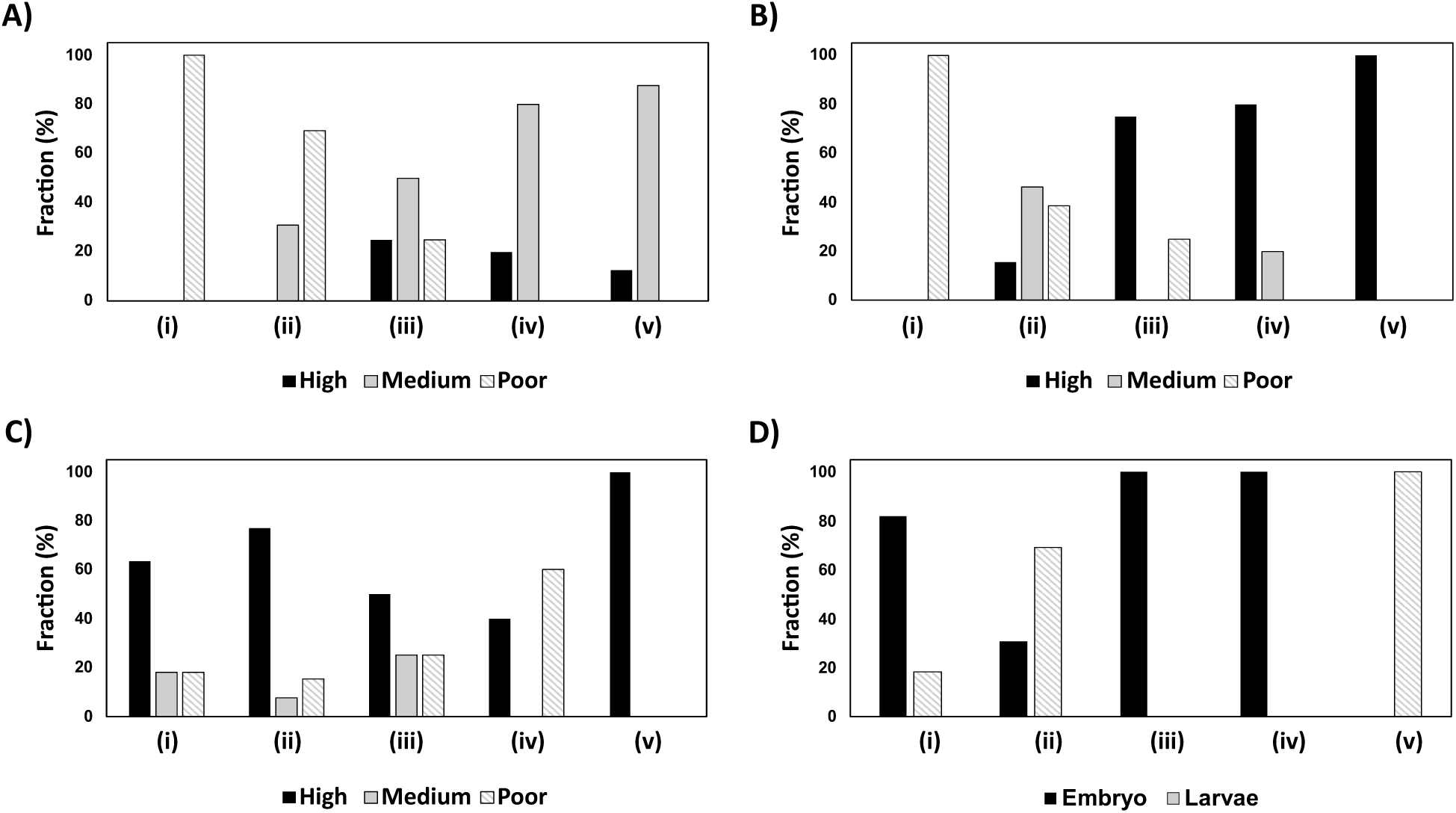
Qualitative assessment of peptide **(A)** adsorption on the outer membrane and **(B)** permeation in the interior, together with **(C)** viability and **(D)** developmental stage of dechorionated and dewaxed *Drosophila melanogaster* hosts incubated with either *(i)* no peptide (PBS, pH 7.4), *(ii)* GEGEGEG, *(iii)* RRRRRRRRR, *(iv)* MRKRHASRREK, or *(v)* cyclo[*Glut*-MRKRHASRRE-*K*\*] peptides. Peptide adsorption onto the membrane was assessed as a thin layer of red fluorescence, while peptide permeation in the interior was assessed as diffuse or punctate red fluorescence within the developing embryo or larva. The level of superficial and inner fluorescence (**A** and **B**), as well as host viability **(C)** were qualitatively categorized as either “high”, “medium”, or “poor”. Accordingly, the axis “fraction (%)” indicates the fraction of the hosts tested under conditions *(i) – (v)* that belonged to either the “high”, “medium”, or “poor” categories.

As anticipated, peptide permeation increased as a result of the loss of the wax layer, whose molecular density and strong hydrophobic character oppose a strong barrier to foreign compounds. The selected cyclic peptide cyclo[*Glut*-MRKRHASRRE-*K*\*], in fact, was found to permeate efficiently all the hosts; likewise, a notable increase in permeation was observed with both RRRRRRRRR and MRKRHASRREK (**Figure 3A** and **3B**).

A loss of viability was observed among the dechorionated-and-dewaxed embryos treated with these two peptides. However, as noted above, this cannot be directly imputed to an inherent toxicity of these peptides, since RRRRRRRRR is known to be cyto-compatible^44^ and a loss of viability was observed in 40% of the embryos treated with no peptide (**Figure 3C**); the absence of a causal connection between peptide permeation and toxicity is corroborated by cyclo[*Glut*-MRKRHASRRE-*K*\*], which was shown to permeate all the tested hosts without affecting their viability. Somewhat unexpectedly, all the dechorionated-and-dewaxed embryos incubated with the cyclic peptide transitioned to the larval state within 12 hours (**Figure 3D**). This, however, should not prompt the conclusion that cyclo[*Glut*-MRKRHASRRE-*K*\*] accelerates embryos development, since the dechorionated-only embryos that were permeated by the peptides did not transition to larvae (**Figure 1D**); furthermore, 70% of the embryos incubated with GEGEGEG, despite being only poorly permeated, transitioned to the larval state.

Confocal fluorescence microscopy images of dechorionated-and-dewaxed embryos and larvae confirmed the superior permeating activity and nuclear co-localization of the lead peptide cyclo[*Glut*-MRKRHASRRE-*K*\*]-K(5-TAMRA) (**Figure 4A**). As noted above, all the hosts incubated with MRKRHASRREK or RRRRRRRRR remained at the embryo stage, whereas those exposed to cyclo[*Glut*-MRKRHASRRE-*K*\*]-K(5-TAMRA) progressed to the larval stage, as evident in **Figure 4A**. The permeation of the linear precursor TAMRA-MRKRHASRREK and the positive control TAMRA-RRRRRRRRR is exemplified in **Figure 4B** and **4C**, respectively: the former produced a diffused fluorescence throughout the embryos, with no defined overlap with the nuclear staining, whereas the latter featured a lesser permeation activity, producing a halo-like pattern around the periphery of the embryo. Finally, the poor permeation of GEGEGEG (**Figure 4D**) is evidenced by the comparison with the embryos that were not exposed to any peptide (**Figure 4E**). Collectively, these results suggest that, while upon removal of the wax layer the embryos become more permeable, a peptide that is specifically designed as an embryo-permeating vector is needed to achieve efficient access to the hosts.

**Figure 4.**
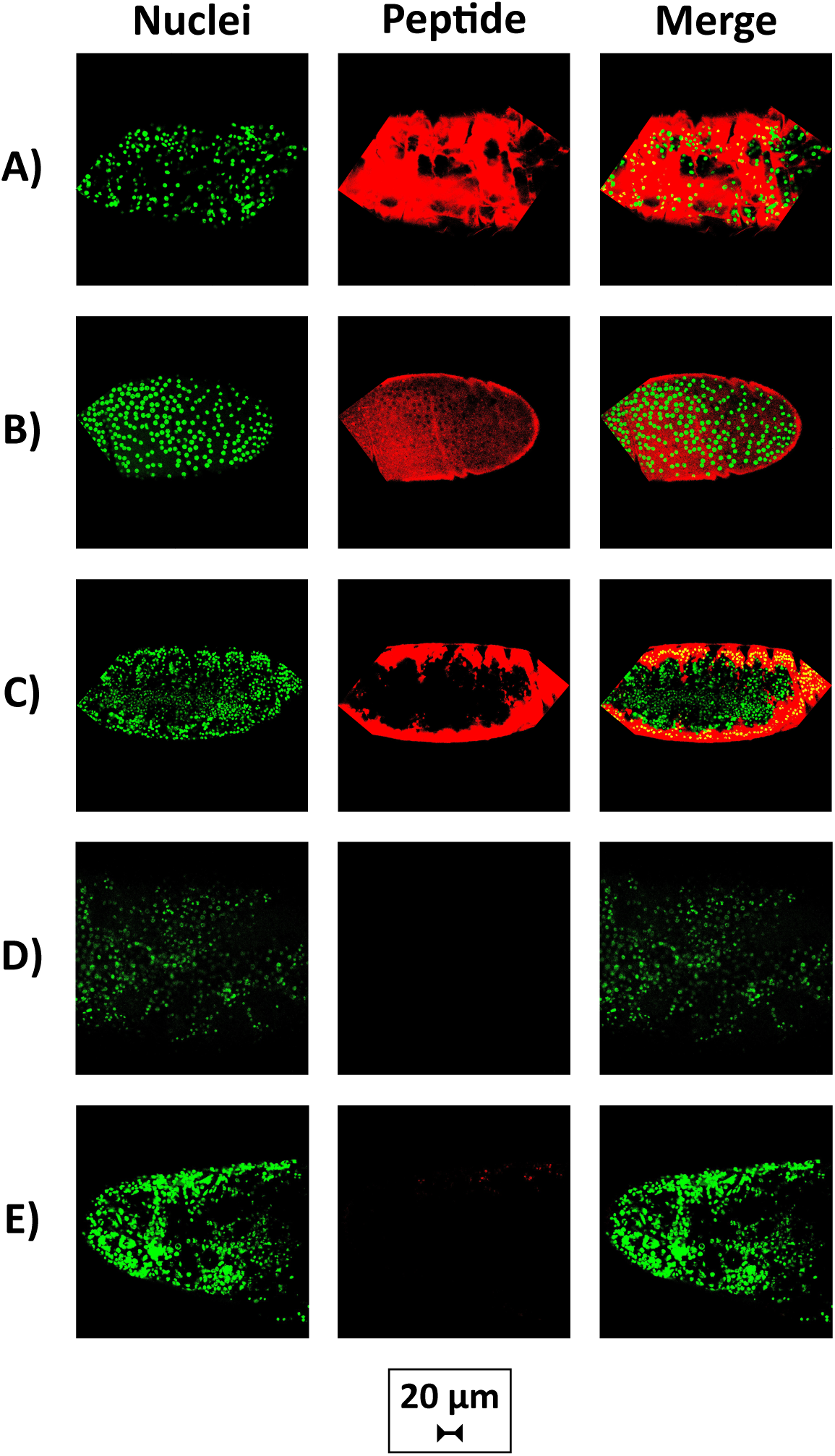
Confocal fluorescence microscopy images of dechorionated and dewaxed *D. melanogaster* embryos incubated with TRITC-labeled peptides **(A)** cyclo[*Glut*-MRKRHASRRE-*K*\*], **(B)** MRKRHASRREK, **(C)** RRRRRRRRR, **(D)** GEGEGEG at 20 μM, or **(E)** no peptide (PBS, pH 7.4), overnight at room temperature (*note:* the scalebar is reported separately from the microscopy images to improve its legibility). Nuclear GFP expression is shown in the green channel, and distribution of permeated peptides in the hosts is shown in the red channel.

### Permeability of cyclo[Glut-MRKRHASRRE-K*] in NIH 3T3 mouse embryonic fibroblast cells

Having demonstrated the permeation of cyclo[*Glut*-MRKRHASRRE-*K*\*]-K(5-TAMRA) in fruit fly embryos, we sought to determine its ability to permeate cell lines that are relevant models in developmental biology. While rich in cationic residues, like many well-known CPPs, the peptide lacks the amphiphilic character (*i*.*e*., the combination of polar and hydrophobic residues) typical of the pore-forming CPPs, which can cause severe cell damage.^45^ We therefore evaluated the permeability of cyclo[*Glut*-MRKRHASRRE-*K*\*], its linear cognate MRKRHASRREK, and the positive and negative controls RRRRRRRRR and GEGEGEG into NIH 3T3 mouse embryonic fibroblast cell line.

The extent of cell penetration of each peptide was measured by flow cytometry single cell analysis of NIH 3T3 cells following incubation with the fluorescently labeled peptides under physiology cell culture conditions. The mean fluorescence intensity (MFI) values of the cells incubated with the various peptides (**Figure 5A**), derived by averaging the flow cytometry fluorescence data (**Figure 5B and 5C**), demonstrate that cyclo[*Glut*-MRKRHASRRE-*K*\*] possesses a superior cell penetration power compared to its linear precursor and the control peptides. The effect of incubation temperature and peptide dose on the resulting cell fluorescence was also evaluated to gain insight into to the mechanism of penetration (active endocytosis *vs*. passive diffusion). A statistically significant difference in cell permeation was observed with cyclo[*Glut*-MRKRHASRRE-*K*\*]-K(5-TAMRA) by increasing the incubation temperature from 4°C, where the peptides access the cells solely by passive diffusion, to physiological conditions (37°C), where the mechanism of active endocytosis become relevant (**Figure 5A**). On the other hand, no statistically significant difference in cell penetration with incubation temperature was observed between the linear precursor and the control peptides. Therefore, as observed with embryo permeation, peptide cyclization appears to play a critical role in endowing the selected sequence with cell-penetrating activity. Furthermore, **Figure 5D** shows an increase in cell permeation with peptide concentration in solution (from 1 to 5 µM). We finally evaluated the safety of cyclo[*Glut*-MRKRHASRRE-*K*\*]-K(5-TAMRA) by measuring the viability of NIH 3T3 cells incubated with the peptide at high concentration (10 µM) for 24 and 48 hrs. The percentage of viable cells shown in **Figure 5E** demonstrates that there is no significant difference on cell viability after incubation with the cyclo[*Glut*-MRKRHASRRE-*K*\*]-K(5-TAMRA) peptide compared to the other control peptides.

**Figure 5.**
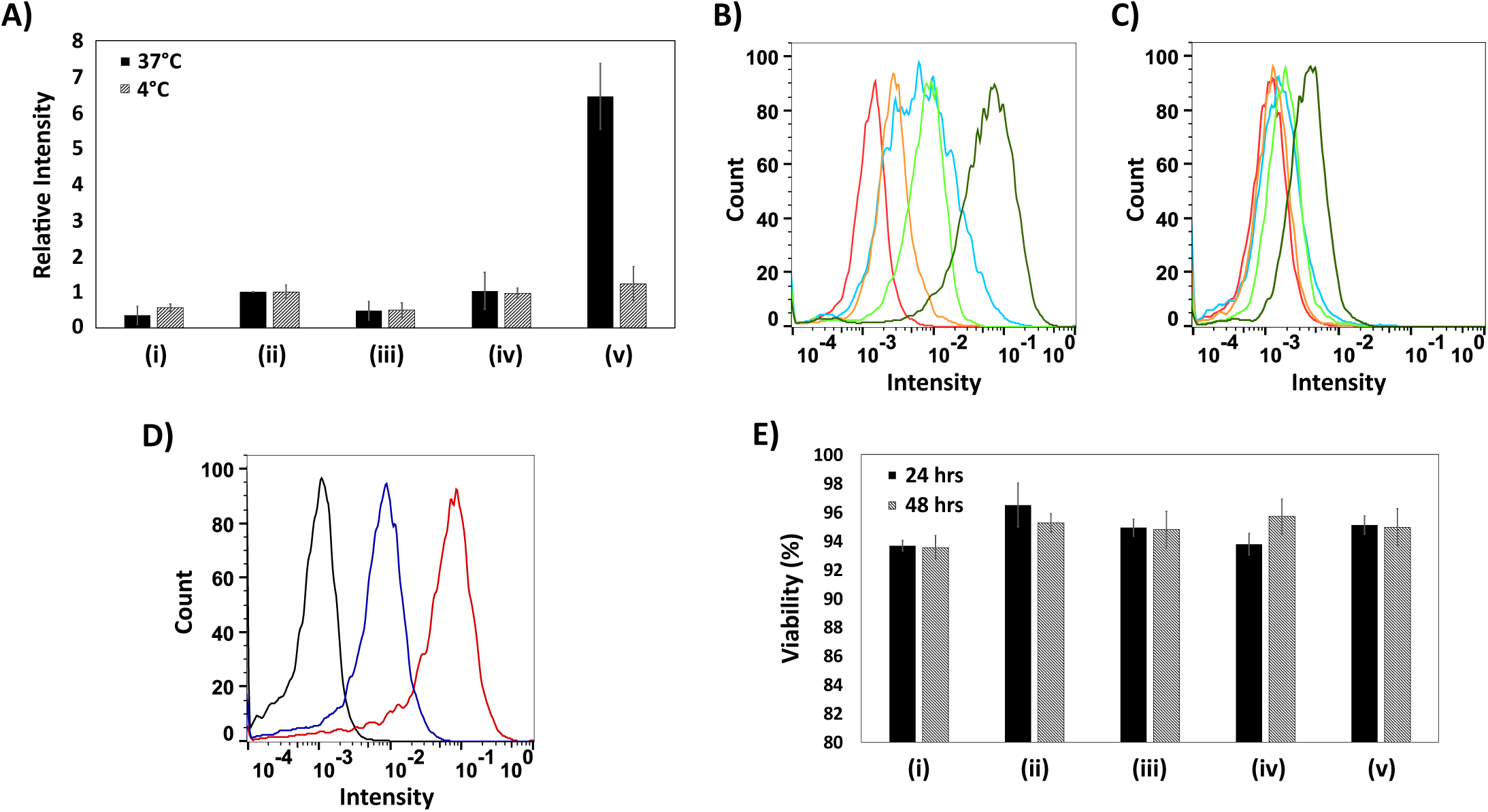
NIH 3T3 peptide permeation comparison with flow cytometry. **(A)** Mean fluorescence intensity (MFI) data for unlabeled cells, cells incubated with TAMRA-labeled peptides: *(i)* no peptide, *(ii)* RRRRRRRRR, *(iii)* GEGEGEG, *(iv)* MRKRHASRREK, or *(v)* cyclo[*Glut*-MRKRHASRRE-*K*\*]; the values of MFI are normalized against those obtained with TAMRA-RRRRRRRRR at the corresponding temperature (*i*.*e*., 4°C or 37°C). **(B)** Representative flow cytometry histogram at 37°C of unlabeled cells (red), cells incubated with TAMRA-labeled peptides: 2 μM RRRRRRRRR (cyan), 2 μM GEGEGEG (orange), 2 μM MRKRHASRREK (light green), or 2 μM cyclo[*Glut*-MRKRHASRRE-*K*\*] (dark green). **(C)** Representative flow cytometry histogram at 4°C of unlabeled cells (red), cells incubated with TAMRA-labeled peptides: 2 μM RRRRRRRRR (cyan), 2 μM GEGEGEG (orange), 2 μM MRKRHASRREK (light green), or 2 μM cyclo[*Glut*-MRKRHASRRE-*K*\*] (dark green). **(D**) Representative flow cytometry analysis of NIH 3T3 cells incubated with 1 μM (blue) or 5 μM (red) of cyclo[*Glut*-MRKRHASRRE-*K*\*]in DMEM, 10% FBS overnight at 37°C. Cells alone are shown in black. **(E)** Percentage of viable cells after 24 or 48 hour incubation with *(i)* PBS or *(ii) - (v)* 10 μM of each peptide. Data analysis using a two-tailed t-test assuming unequal variances indicates a statistical higher permeation of M-cyclo[*Glut*-MRKRHASRRE-*K*\*] compared to the linear precursor and the control peptides at 37°C (p<0.05). Error bars correspond to the standard error from 3 independent repeats.

The intracellular trafficking of the peptide in NIH 3T3 cells was tracked via laser-scanning confocal fluorescence microscopy (**Figure 6**); because cell fixation can cause an artificial increase in fluorescence^46–48^, live cell images were obtained. Cell imaging confirms that cyclo[*Glut*-MRKRHASRRE-*K*\*] affords a substantially greater permeation than its linear cognate MRKRHASRREK and the positive control RRRRRRRRR, as indicated by the diffuse and punctate fluorescence in **Figure 6A** in contrast with the spare punctuate fluorescence in **Figures 6B** and **6C**; as anticipated, no permeation was detected with the negative control peptide GEGEGEG (**Figure 6D**).

**Figure 6.**
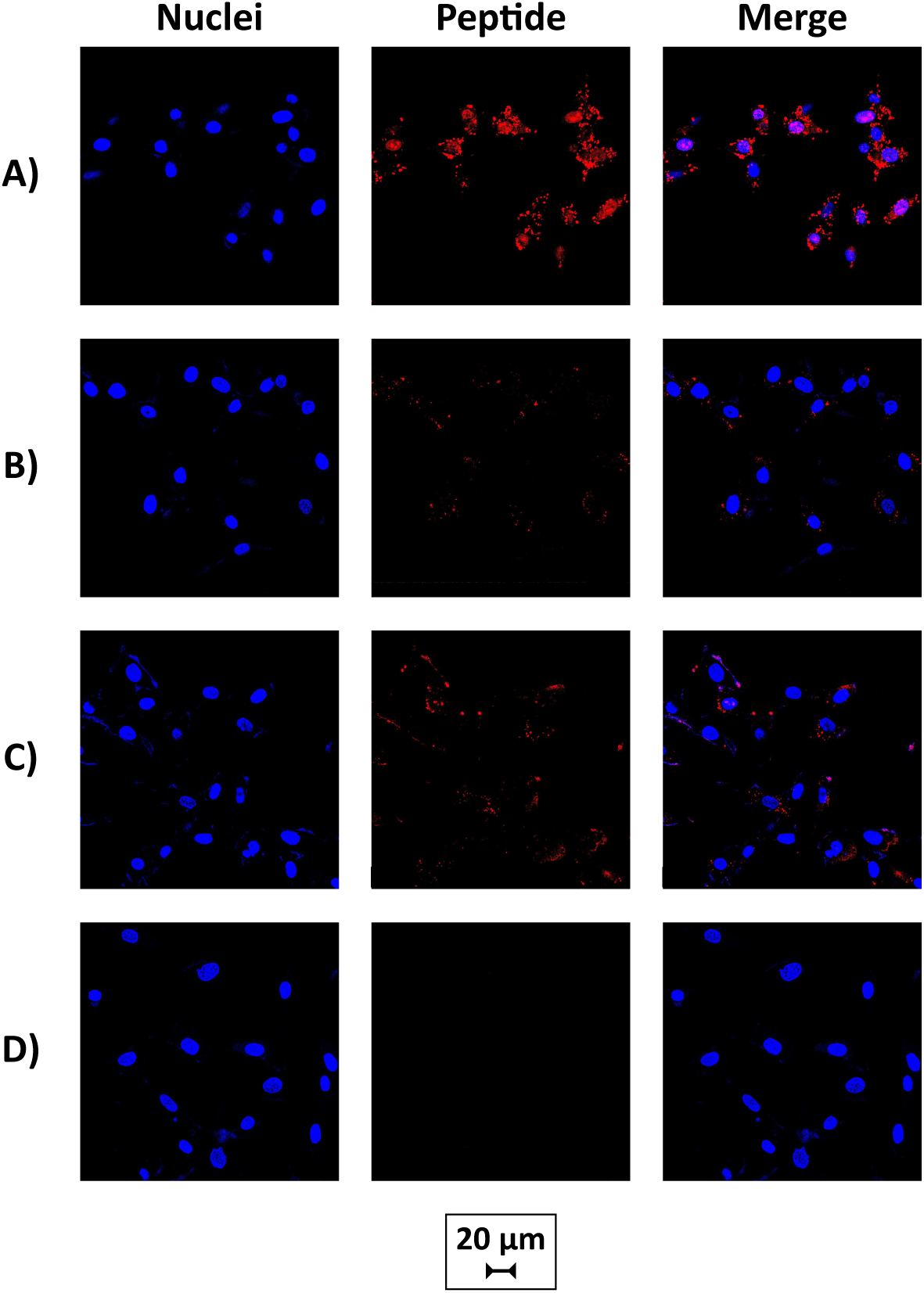
Confocal microscopy images of NIH 3T3 cells incubated with 5 μM TAMRA-labeled peptide in Opti-mem media supplemented with 2% Fetal Bovine Serum for 3 hours at 37°C: **(A)** of cyclo[*Glut*-MRKRHASRRE-*K*\*]; **(B)** 5 μM of MRKRHASRREK; **(C)** 5 μM of RRRRRRRRR; or **(D)** 5 μM of GEGEGEG. Blue channel indicates nuclei staining with Hoecht and red channel indicates the peptide. *Note:* the scalebar is reported separately from the microscopy images to improve its legibility.

### Permeability of cyclo[Glut-MRKRHASRRE-K*] in H9 human embryonic stem cells (hESCs)

The cell permeation assays outlined above were repeated with H9 human embryonic stem cells (hESCs). Unlike NIH 3T3 cells, H9 cells grow in colonies and require a different basal growth media. Notably, penetration of other pluripotent stem cells using RRRRRRRRR and the HIV cell-penetrating peptide TAT (GRKKRRQRRRPQ) has been demonstrated in the literature.^32,33^ To our knowledge, however, no stem cell-penetrating peptides have been discovered using a *de novo* combinatorial approach like the one presented here.

Cyclo[*Glut*-MRKRHASRRE-*K*\*] and its linear counterpart showed comparable permeation, although both were outperformed by the positive control peptide RRRRRRRRR (**Figure 7A**). Interestingly, no statistically significant difference based on temperature was observed in the comparison among the various peptides; however, the comparison with the control peptides at different temperatures (4°C *vs*. 37°C), obtained by averaging the flow cytometry histograms (**Figure 7B** and **7C**), shows a slightly different peptide permeation behavior compared to that observed with 3T3 cells. Interestingly, a bimodal histogram was observed with RRRRRRRRR incubated hESC cells (**Figure 7B**), which could be attributed to differences in either colony or single cell permeation (this phenomenon was not explored further). However, it must be emphasized that data from these flow cytometry studies need to be interpreted with caution, and in conjunction with microscopy studies (**Figure 8**), since non-specific binding of peptides to the cell surface may occur. Finally, the biocompatibility of cyclo[*Glut*-MRKRHASRRE-*K*\*]-K(5-TAMRA) on H9 cells was evaluated, indicating no significant impact on cell viability after incubation with the cyclo[*Glut*-MRKRHASRRE-*K*\*]-K(5-TAMRA) peptide (**Figure 7E**).

**Figure 7.**
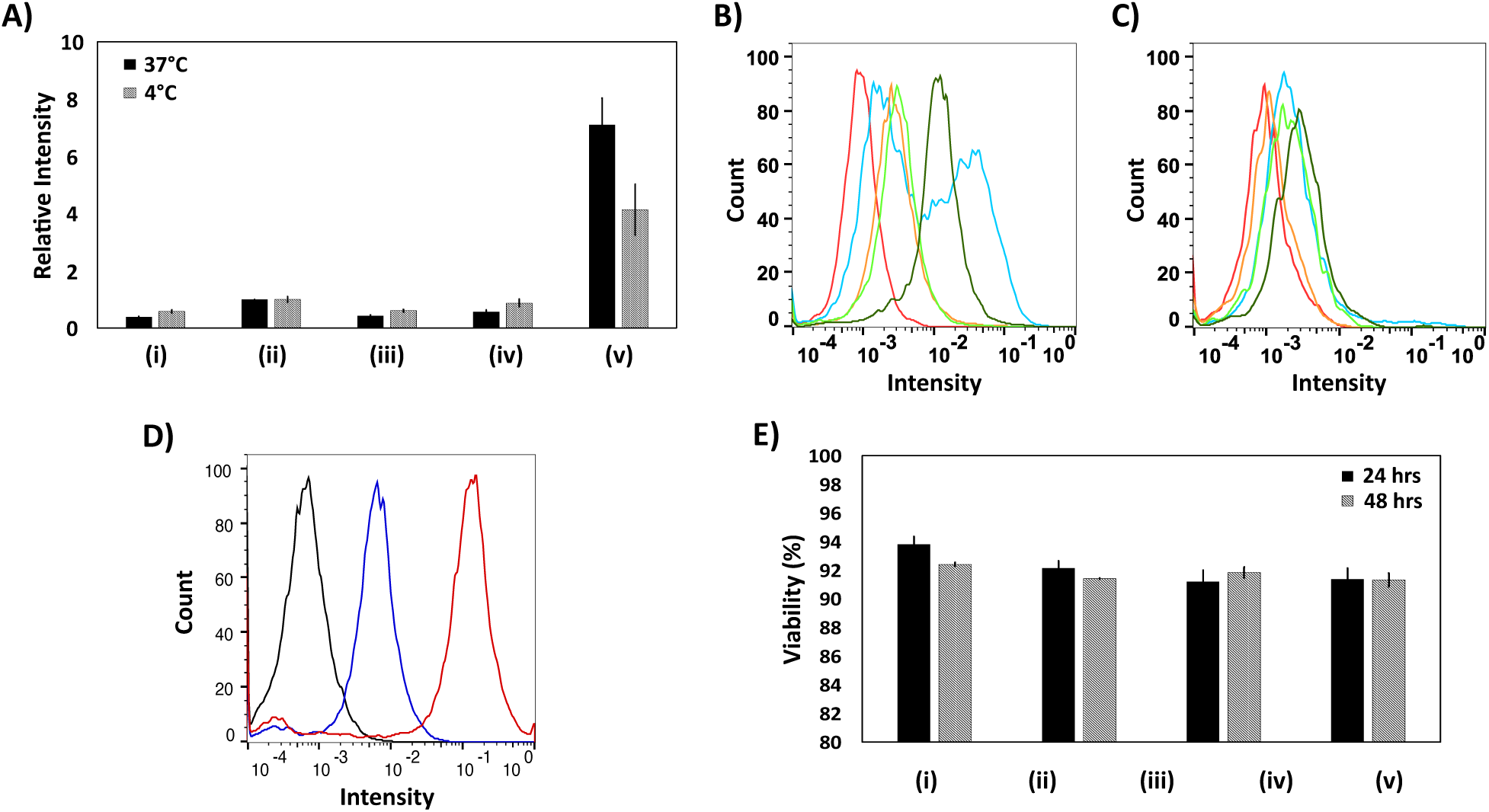
H9 human embryonic stem cell peptide permeation comparison with flow cytometry. **(A)** Mean fluorescence intensity (MFI) data for unlabeled cells, cells incubated with TAMRA-labeled peptides: *(i)* no peptide, *(ii)* RRRRRRRRR, *(iii)* GEGEGEG, *(iv)* MRKRHASRREK, or *(v)* cyclo[*Glut*-MRKRHASRRE-*K*\*]; the values of MFI are normalized against those obtained with TAMRA-RRRRRRRRR at the corresponding temperature (*i*.*e*., 4°C or 37°C). **(B)** Representative flow cytometry histogram at 37°C of unlabeled cells (red), cells incubated with TAMRA-labeled peptides: 2 μM RRRRRRRRR (cyan), 2 μM GEGEGEG (orange), 2 μM MRKRHASRREK (light green), or 2 μM cyclo[*Glut*-MRKRHASRRE-*K*\*] (dark green). **(C)** Representative flow cytometry histogram at 4°C of unlabeled cells (red), cells incubated with TAMRA-labeled peptides: 2 μM RRRRRRRRR (cyan), 2 μM GEGEGEG (orange), 2 μM MRKRHASRREK (light green), or 2 μM cyclo[*Glut*-MRKRHASRRE-*K*\*] (dark green). **(D**) Representative flow cytometry analysis of H9 hESCs incubated with 1 μM (blue) or 5 μM (red) of cyclo[*Glut*-MRKRHASRRE-*K*\*] in E8 media overnight at 37°C. Cells alone are shown in black. **(E)** Percentage of viable cells after 24 or 48 hour incubation with *(i)* PBS or *(ii) - (v)* 10 μM of each peptide. Data analysis using a two-tailed t-test assuming unequal variances indicates a statistical higher permeation of RRRRRRRRR compared to all other peptides at 37°C (p<0.05). No statistically significant differences in viability were observed. Error bars correspond to the standard error from 3 independent repeats.

**Figure 8.**
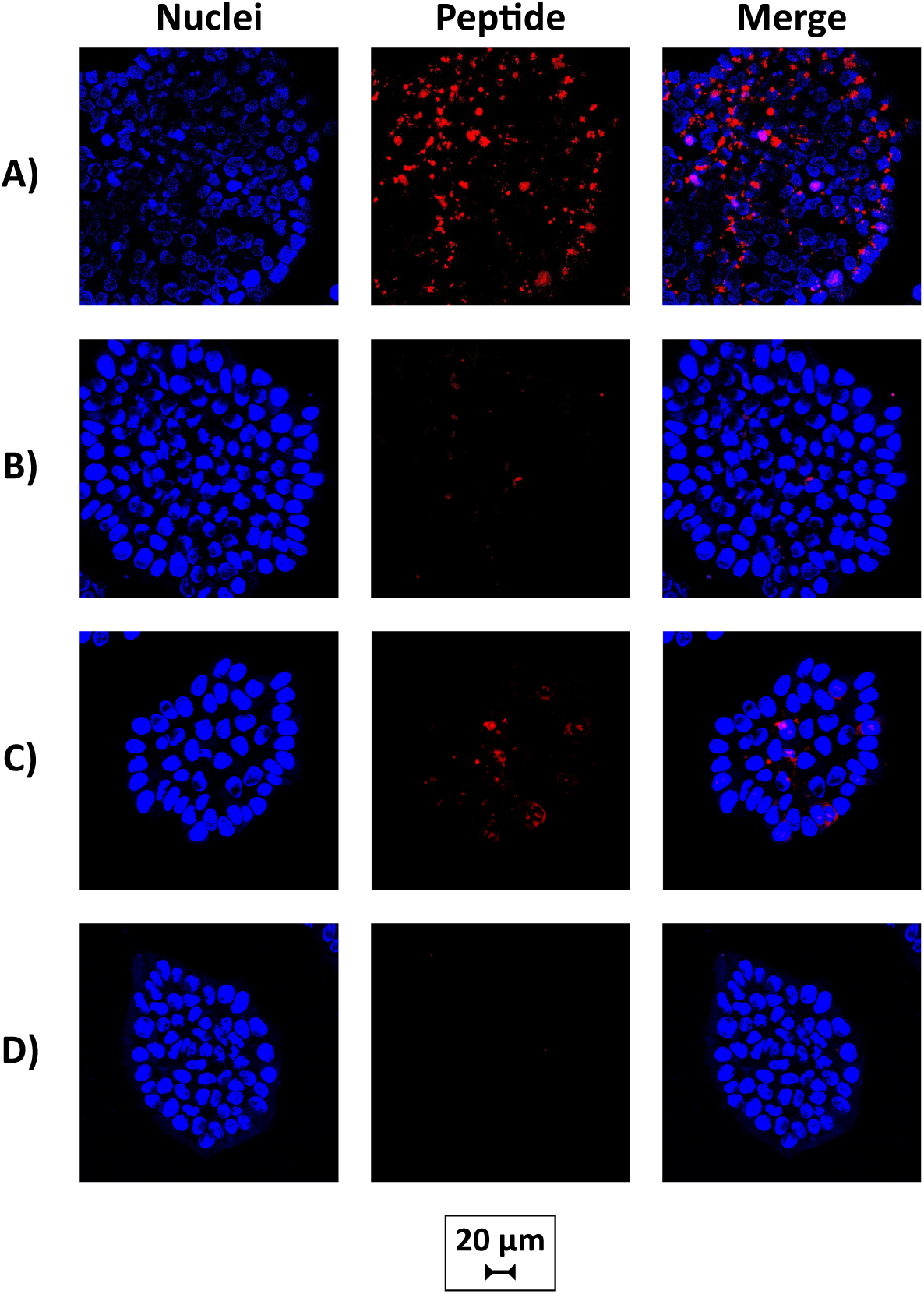
Confocal microscopy images of H9 human embryonic stem cells incubated with 1 μM TAMRA-labeled peptide in E8 media for 3 hours at 37°C: **(A)** of cyclo[*Glut*-MRKRHASRRE-*K*\*]; **(B)** 5 μM of MRKRHASRREK; **(C)** 5 μM of RRRRRRRRR; or **(D)** 5 μM of GEGEGEG. Blue channel indicates nuclei staining with Hoecht and red channel indicates the peptide. *Note:* the scalebar is reported separately from the microscopy images to improve its legibility.

Confocal fluorescence microscopy images of hESC colonies incubated with labeled peptides indicate that cyclo[*Glut*-MRKRHASRRE-*K*\*] (**Figure 8A**) produced a slightly higher fluorescence than RRRRRRRRR (**Figure 8C**). It is also noted that cyclo[*Glut*-MRKRHASRRE-*K*\*] penetration resulted in a punctuate-only fluorescence in hESC colonies, in contrast with what observed with 3T3 cells, where a diffuse and punctuate fluorescence was obtained. This suggests a potential sequestration of the cyclic peptides within endosomes or lysosomes upon access into hESC colony. As anticipated, neither the linear cognate MRKRHASRREK nor the negative control GEGEGEG afforded any significant penetration (**Figure 8B** and **8D**) with a weak punctate fluorescence occurring for the linear peptide only. These data indicate that cyclo[*Glut*-MRKRHASRRE-*K*\*] can penetrate H9 hESCs. Collectively, these data demonstrate that cyclo[*Glut*-MRKRHASRRE-*K*\*] is capable of permeating mammalian stem cells rapidly and efficiency.

## Conclusions

We have successfully developed a novel screening technique that uses mRNA display libraries of cyclic peptides to identify *de novo* peptide capable of *Drosophila melanogaster* permeation. To our knowledge, this is the first peptide identified to permeate into *Drosophila melanogaster*. We further characterized cyclo[*Glut*-MRKRHASRRE-*K*\*]-K(5-TAMRA) on two other cell types to show its utility as a general membrane-permeating peptide. In all cases, the peptide was able to penetrate into the cell or organism of interest. Also in all cases, the cyclization of the peptide was necessary to enhance its permeability. Membrane-penetrating peptides developed using this approach may have application as a delivery agents, which can supplement or displace traditional delivery methods such as microinjections or liposomal-based delivery methods. Our results demonstrate the usefulness of our approach for rapid identification of cyclic MPPs.

## Acknowledgements

This work was funded by the National Science Foundation (CBET 1510845). JB kindly acknowledge support from a National Institute of Health Molecular Biotechnology Training Fellowship (NIH T32 GM008776). We would also like to thank the UNC High-Throughput Peptide Synthesis and Array facility for peptide synthesis.

## Conflict of interest statement

The authors do not have any conflict of interest to acknowledge.

## Notes

### Competing Interest Statement

The authors have declared no competing interest.

## References

1. Sonane, M., Goyal, R., Chowdhuri, D. K., Ram, K. R. & Gupta, K. C. Enhanced efficiency of P-element mediated transgenesis in Drosophila : Microinjection of DNA complexed with nanomaterial. Sci. Rep. 3, 3408 (2013).

2. Zabihihesari, A., Hilliker, A. J. & Rezai, P. Localized microinjection of intact Drosophila melanogaster larva to investigate the effect of serotinin on heart rate. Lab Chip 20, 343–355 (2020).

3. Szelei, J. et al. Liposome-mediated gene transfer in fish embryos. Transgenic Res. 3, 116–119 (1994).

4. Wagstaff, K. & Jans, D. Protein Transduction: Cell Penetrating Peptides and Their Therapeutic Applications. Curr. Med. Chem. 13, 1371–1387 (2006).

5. Lakshmanan, M., Kodama, Y., Yoshizumi, T., Sudesh, K. & Numata, K. Rapid and Efficient Gene Delivery into Plant Cells Using Designed Peptide Carriers. Biomacromolecules 14, 10–16 (2013).

6. Joo, J. Y. et al. VisuFect-mediated siRNA delivery into zygotes. Colloids Surfaces B Biointerfaces 135, 646–651 (2015).

7. Liu, Y. et al. Enhancing gene delivery of adeno-associated viruses by cell-permeable peptides. Mol. Ther. - Methods Clin. Dev. 1, 12 (2014).

8. Gaston, J. et al. Intracellular delivery of therapeutic antibodies into specific cells using antibody-peptide fusions. Sci. Rep. 9, 18688 (2019).

9. Trabulo, S., Cardoso, A. L., Mano, M. & Pedroso de Lima, M. C. Cell-Penetrating Peptides—Mechanisms of Cellular Uptake and Generation of Delivery Systems. Pharmaceuticals 3, 961–993 (2010).

10. Lee, J. H., Song, H. S., Lee, S. G., Park, T. H. & Kim, B. G. Screening of cell-penetrating peptides using mRNA display. Biotechnol. J. 7, 387–396 (2012).

11. Agrawal, P. et al. CPPsite 2. 0 : a repository of experimentally validated cell-penetrating peptides. Nucleic Acids Res. 44, 1098–1103 (2016).

12. Milletti, F. Cell-penetrating peptides: Classes, origin, and current landscape. Drug Discov. Today 17, 850–860 (2012).

13. Derakhshankhah, H. & Jarari, S. Cell penetrating peptides : A concise review with emphasis on biomedical applications. Biomed. Pharmacother. 108, 1090–1096 (2018).

14. Wang, H. & Liu, R. Advantages of mRNA display selections over other selection techniques for investigation of protein-protein interactions. Expert Rev. Proteomics 8, 335–346 (2011).

15. Hoffmann, K. et al. A platform for discovery of functional cell-penetrating peptides for efficient multi-cargo intracellular delivery. Sci. Rep. 8, 12538 (2018).

16. Qian, Z. et al. Efficient Delivery of Cyclic Peptides into Mammalian Cells with Short Sequence Motifs. ACS Chem. Biol. 8, 423–431 (2012).

17. Qian, Z. et al. Discovery and Mechanism of Highly Efficient Cyclic Cell-Penetrating Peptides. Biochemistry 55, 2601–2612 (2016).

18. Qian, Z. et al. Early Endosomal Escape of a Cyclic Cell-Penetrating Peptide Allows Effective Cytosolic Cargo Delivery. Biochemistry 53, 4034–4046 (2014).

19. Lättig-Tünnemann, G. et al. Backbone rigidity and static presentation of guanidinium groups increases cellular uptake of arginine-rich cell-penetrating peptides. Nat. Commun. 2, 453 (2011).

20. Mandal, D., Shirazi, A. N. & Parang, K. Cell-Penetrating Homochiral Cyclic Peptides as Nuclear-Targeting Molecular Transporters. Angew Chem Int Ed Engl. 50, 9633–9637 (2011).

21. Traboulsi, H. et al. Macrocyclic Cell Penetrating Peptides: A Study of Structure-Penetration Properties. Bioconjug. Chem. 26, 405–411 (2015).

22. Menegatti, S., Hussain, M., Naik, A. D., Carbonell, R. G. & Rao, B. M. mRNA Display Selection and Solid-Phase Synthesis of Fc-Binding Cyclic Peptide Affinity Ligands. Biotechnol. Bioeng. 110, 857–870 (2013).

23. Rand, M. D., Kearney, A. L., Dao, J. & Clason, T. Permeabilization of Drosophila embryos for introduction of small molecules. Insect Biochem. Mol. Biol. 40, 792–804 (2010).

24. Papassideri, I., Margaritis, L. H. & Gulik-Krzywicki, T. The Egg-Shell of Drosophila Melanogaster VI, Structural Analysis of the Wax layer in Laid Eggs. Tissue Cell 23, 567–575 (1991).

25. Margaritis, L. H., Kafatos, F. C. & Petrij, W. H. The Eggshell of Drosophila Melanogaster Fine Structure of the Layers and Regions of the Wild-type Eggshell. J. Cell Sci. 43, 1–35 (1980).

26. Lee, J.-S. et al. RNA-Guided Genome Editing in Drosophila with the Purified Cas9 Protein. G3 Genes, Genomes, Genet. 4, 1291–1295 (2014).

27. Heigwer, F., Port, F. & Boutros, M. RNA Interference (RNAi) Screening in Drosophila. Genetics 208, 853–874 (2018).

28. Schulman, V. K., Folker, E. S. & Baylies, M. K. A method for reversible drug delivery to internal tissues of Drosophila embryos. Fly 7, 193–203 (2013).

29. Chugh, A. & Eudes, F. Study of uptake of cell penetrating peptides and their cargoes in permeabilized wheat immature embryos. FEBS J. 275, 2403–2414 (2008).

30. Joo, J. Y. et al. Microinjection free delivery of miRNA inhibitor into zygotes. Sci. Rep. 4, 5417 (2014).

31. Yang, N. J. & Hinner, M. J. Getting Across the Cell Membrane: An Overview for Small Molecules, Peptides, and Proteins. Methods Mol. Biol. 1266, 29–53 (2015).

32. Kaitsuka, T. & Tomizawa, K. Cell-Penetrating Peptide as a Means of Directing the Differentiation of Induced Pluripotent Stem Cells. Int. J. Mol. Sci. 16, 26667–26676 (2015).

33. Yukawa, H. et al. Transduction of cell-penetrating peptides into induced pluripotent stem cells. Cell Transplant. 19, 901–909 (2010).

34. Ng, K. K. et al. Intracellular Delivery of Proteins via Fusion Peptides in Intact Plants. PLoS One 11, e0154081 (2016).

35. Frankel, A. D. & Pabo, C. O. Cellular Uptake of the Tat Protein from Human lmmunodeficiency Virus. Cell 55, 1189–1193 (1988).

36. Green, M. & Loewenstein, P. M. Autonomous Functional Domains of Chemically Synthesized Human lmmunodeficiency Virus Tat Trans-Activator Protein. Cell 55, 1179–1188 (1988).

37. Green, M., Ishino, M. & Loewenstein, P. M. Mutational Analysis of HIV-1 Tat Minimal Domain Peptides : Identification of Trans-Dominant Mutants That Suppress HIV-LTR-Driven Gene Expression. Cell 58, 215–223 (1989).

38. Derossi, D., Joliot, A. H., Chassaing, G. & Prochiantz, A. The Third Helix of the Antennapedia Homeodornain Translocates through Biological Membranes. J. Biol. Chem. 269, 10444–10450 (1994).

39. Bedewy, W. et al. Generation of a cell-permeable cycloheptapeptidyl inhibitor against the peptidyl-prolyl isomerase Pin1. Org. Biomol. Chem. 15, 4540–4543 (2017).

40. Menegatti, S. et al. De Novo Design of Skin-Penetrating Peptides for Enhanced Transdermal Delivery of Peptide Drugs. Adv. Healthc. Mater. 5, 602–609 (2016).

41. Milletti, F. Cell-penetrating peptides : classes, origin, and current landscape. Drug Discov. Today 17, 850–860 (2012).

42. Wu, T., Manogaran, A. L., Beauchamp, J. M. & Waring, G. L. Drosophila vitelline membrane assembly: A critical role for an evolutionarily conserved cysteine in the ‘VM domain’ of sV23. Dev. Biol. 347, 360–368 (2010).

43. Petrij, W. H., Wyman, A. R. & Kafatos, F. C. Specific Protein Synthesis in Cellular Differentiation III. The Eggshell Proteins of Drosophila melanogaster and Their Program of Synthesis. Dev. Biol. 49, 185–199 (1976).

44. Kanekura, K. et al. Characterization of membrane penetration and cytotoxicity of C9orf72-encoding arginine-rich dipeptides. Sci. Rep. 12740 (2018).

45. Kilk, K., Mahlapuu, R., Soomets, U. & Langel, Ü. Analysis of in vitro toxicity of five cell-penetrating peptides by metabolic profiling. Toxicology 265, 87–95 (2009).

46. Richard, J. P. et al. Cell-penetrating Peptides - A Reevaluation of the mechanism of cellular uptake. J. Biol. Chem. 278, 585–590 (2003).

47. Mano, M., Teodosio, C., Paiva, A., Simoes, S. & Pedroso de Lima, M. C. On the mechanisms of the internalization of S413-PV cell-penetrating peptide. Biochem. J. 390, 603–612 (2005).

48. Lundberg, M., Wikstrom, S. & Johansson, M. Cell Surface Adherence and Endocytosis of Protein Transduction Domains. Mol. Ther. 8, 143–150 (2003).

