## Supporting information for "Discovery of membrane-permeating cyclic peptides via mRNA display"

**Table S1.** Oligonucleotides encoding randomized peptides

| | Sequence |
|---|---|
| Oligo 1 | 5'-GGA CAA TTA CTA TTT ACA ATT ACA ATG NNN NNN NNN NNN NNN NNN NNN NNN NNN NNN AAA GGC AGC GGC TCC GGT CAT CAC CAC CAT CAC CAT ATG GGA ATG-3' |

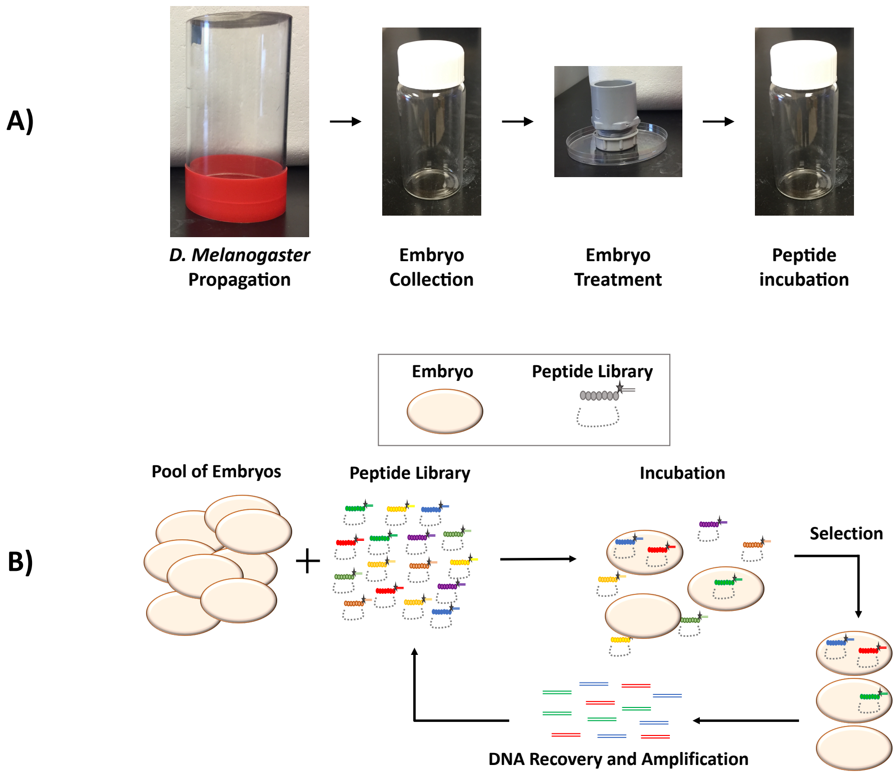


**Figure S1.** *Drosophila melanogaster* embryo handling and mRNA display screening procedure. **(A)** Components utilized for *D. melanogaster* growth, embryo laying and development, and subsequent embryo handling: the *D. melanogaster* are initially placed in a cage with grape juice agar plate supplemented with yeast paste and allowed to mate; the embryos are collected, washed, and transferred to a gray collection basket with a mesh screen for dechorionation or dechorionation/dewaxing; the embryos are finally transferred to a scintillation and incubated with either the peptide library or individual embryo-permeating peptides. **(B)** Schematics of the screening procedure: the dechorionated/dewaxed embryos are incubated with the library; following incubation, the embryos are lysed, and the library-associated cDNA is amplified via PCR and utilized to construct subsequent libraries; starting at the second round of screening, a membrane/organelle centrifugation step is added after the lysis of the embryos to avoid the selection of peptide sequences that are absorbed within the embryo vitelline membrane but do not possess a true permeation activity (*i.e.*, false positives).

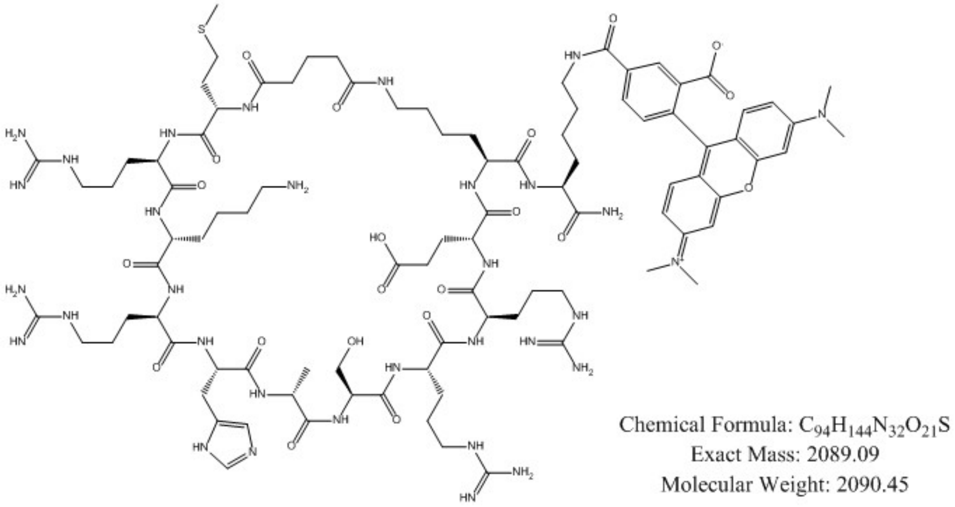


**Figure S2.** Structure and molecular properties of the lead candidate cyclo[*Glut*-MRKRHASRRE-*K**] conjugated to a TAMRA fluorophore.

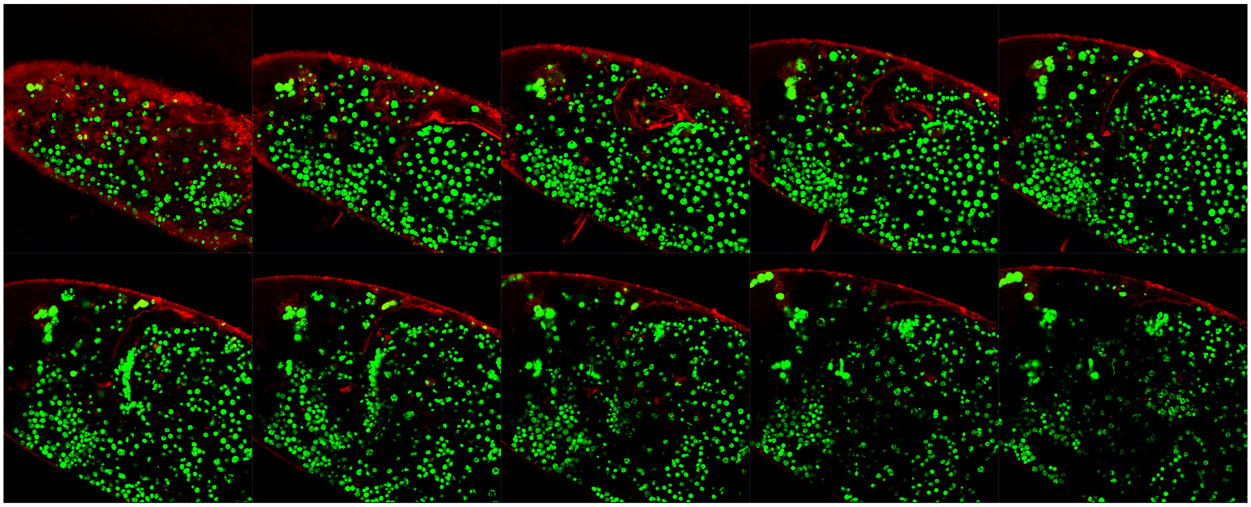

**Figure S3.** Confocal fluorescence representative z-stack microscopy images of dechorionated *D. melanogaster* embryos incubated with cyclo[*Glut*-MRKRHASRRE-*K**] at a concentration of 20 μM, overnight at room temperature. Nuclear GFP expression is shown in the green channel, and distribution of permeated peptides in the embryos is shown in the red channel.
